# Bioregional boundaries and genomically-delineated stocks in snapper (*Chrysophrys auratus*) from southeastern Australia

**DOI:** 10.1101/2023.01.16.524335

**Authors:** Andrea Bertram, Justin Bell, Chris Brauer, Anthony Fowler, Paul Hamer, Jonathan Sandoval-Castillo, John Stewart, Maren Wellenreuther, Luciano B. Beheregaray

## Abstract

Marine species often exhibit genetic discontinuities concordant with biogeographic boundaries, frequently occurring due to changes in ocean circulation, bathymetry, coastline topography and temperature. Here we used 10,916 single nucleotide polymorphisms (SNPs) to assess the concordance between population genomic differentiation and coastal biogeography in the fishery important snapper (*Chrysophrys auratus*) across southeastern Australia. Additionally, we investigated whether spatial scales of assessment and management of snapper align with evidence from population genomics. Across 488 snapper samples from 11 localities between the west coast of South Australia and the south coast of New South Wales, we detected genomic structure concordant with the region’s three biogeographic provinces. We also detected fine-scale genetic structuring relating to spatial variation in spawning and recruitment dynamics, as well as temporal stability in the genomic signal associated with two important spawning grounds. The current management boundaries in the region coincided with either the genetic breaks at bioregional boundaries or with localscale variation. Our study highlights the value of population genomic surveys in species with high dispersal potential for uncovering stock boundaries and demographic variation related to spawning and recruitment. It also illustrates the importance of marine biogeography in shaping population structure in commercial species with high dispersal potential.

## Introduction

Despite the scarcity of obvious physical barriers in the ocean, marine species are often genetically structured into discrete populations that are connected by varying degrees of gene flow, even in species with high dispersal potential (Grummer et al., 2019). In the absence of topographical barriers, genetic structure can form in species with high dispersal potential due to oceanographic features like fronts, eddies and currents (e.g., at convergence or divergence zones), as well as environmental gradients in parameters like salinity and temperature (Banks et al., 2007; Xuereb et al., 2018; Grummer et al., 2019; Vera et al., 2022). Additionally, in coastal marine species, patterns of genetic structure are often a result of historical vicariance that formed during periods of lowered sea level (i.e., during glacial maxima) and persisted even after restoration of habitat continuity following inundation of continental shelf regions (Sinclair et al., 2016; Teske et al., 2017). The aforementioned features are often associated with bioregional boundaries, which are frequently observed to coincide with genetic breaks across multiple taxa with a range of life histories. For example, along the Pacific coast of North America are a number of biogeographic barriers, thought to coincide with upwelling regions and changes in bathymetry and ocean circulation, which correspond to genetic structuring in a range of fish and invertebrate species (Davis et al., 1981; Sivasundar and Palumbi, 2010; Briggs and Bowen, 2012; Silliman, 2019).

Understanding how marine species are structured geographically, as well as the levels of connectivity between discrete populations, is vital for effective management. In exploited marine species, this is because differentiated populations not only have different genetic characteristics (i.e., genetic diversity and effective population sizes), but are also expected to have unique demographic characteristics (i.e., mortality, recruitment, growth rates and abundances), and therefore have the potential to respond differently to fishing pressures (Cadrin et al., 2020). Because of this, an objective of fishery assessment and management is to treat differentiated populations as independent units (i.e., stocks; Cadrin, 2020).

The characterisation of genetic structure and population connectivity in marine species is typically difficult because of their often-great abundances and high dispersal capabilities (Gagnaire et al., 2015; Bernatchez et al., 2017; Grummer et al., 2019). However, advances in DNA sequencing technologies and associated downstream analyses, as well as reductions in operating costs, now allow for the generation of very large datasets of genetic markers. Large datasets of genetic markers (e.g., genome-wide single nucleotide polymorphism, SNP datasets) are more powerful at detecting the subtle genetic structure common in marine species (relative to the smaller datasets previously used in population genetic structure studies, i.e., microsatellite DNA datasets). As a result, the value of genetic methods in addressing important fisheries management issues has increased in recent years. There are numerous examples in the recent literature where large SNP datasets were able to detect levels of genetic structuring in fishery important marine species for which microsatellite datasets previously could not (e.g., Van Wyngaarden et al., 2017; Silliman, 2019; Petrolo et al., 2021; Bertram et al., 2022).

Snapper (*Chrysophrys auratus*) is a coastal sea bream distributed across temperate and sub-tropical Australia and New Zealand (Gomon et al., 2008). The species is a highly fecund, multiple batch broadcast spawner, that forms aggregations to breed when water temperatures are between 15 and 22°C (Parsons et al., 2014). Snapper’s pelagic larval stage lasts for 17-33 days, and maturation occurs between 3 and 7 years of age, depending on location (Parsons et al., 2014; Wakefield et al., 2015). Adult fish are known to live for up to ~60 years of age and reach lengths of ~100 cm (Parsons et al., 2014). Although snapper is an important commercial and recreational fishery resource across its range, southeastern Australia (see Figure 1) is a particular hotspot (Fowler et al., 2021). Between 2010 and 2019, South Australia (SA) generally provided the highest state-based commercial catches of snapper in Australia (although note recent declines, detailed below). Victoria (Vic) on the other hand dominates the country’s recreational snapper catch, bringing significant economic and social benefits to the state (Jalali et al., 2022). In 2011/12 in SA, ~875 t of snapper were captured by the commercial sector, while in Vic in 2006/7, ~600 t of snapper were captured by the recreational sector (Fowler et al., 2021). Consequently, snapper resources in southeast region are of the most valuable in Australia.

**Figure 1.**
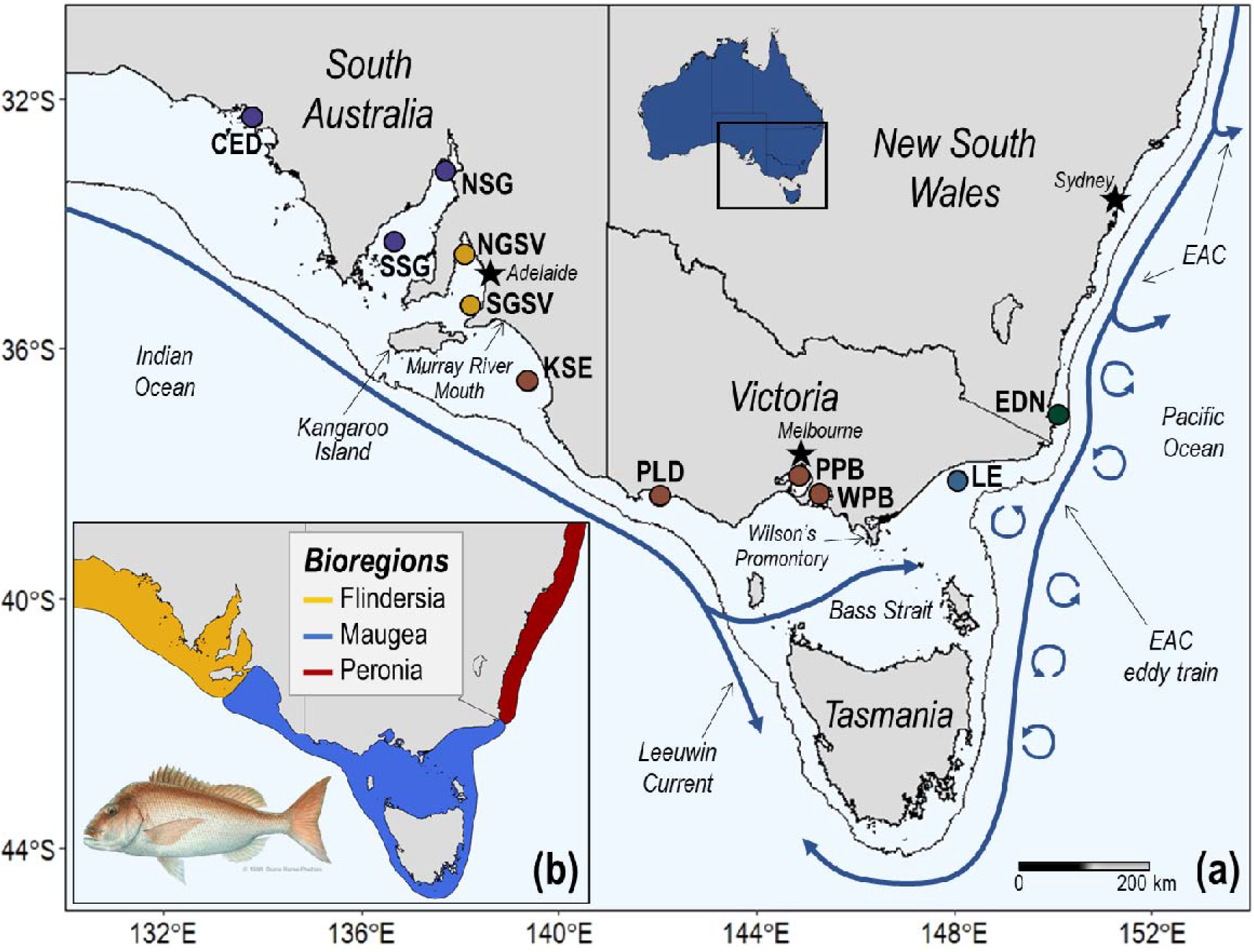
Maps of (a) the sampling sites for snapper (*Chrysophrys auratus*) across southeastern Australia, with sites coloured by the relevant management area: 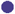 = Spencer Gulf/West Coast, 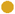 = Gulf St Vincent, 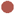 = Western Victoria, 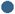 = Eastern Victoria, 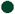 = East Coast; and (b) the major biogeographic provinces in southeastern Australia: Flindersia, Maugea and Peronia. The three stars indicate the capital cities of the relevant southeastern Australian states. CED, Ceduna; NSG, Northern Spencer Gulf; SSG, Southern Spencer Gulf; NGSV, Northern Gulf St Vincent; SGSV, Southern Gulf St Vincent; KSE, Kingston SE; PLD, Portland; PPB, Port Phillip Bay; WPB, Western Port Bay; LE, Lakes Entrance; EDN, Eden.

In southeastern Australia, snapper is currently managed as five separate stocks – Spencer Gulf/West Coast, Gulf St Vincent, Western Victoria, Eastern Victoria and the NSW (New South Wales) component of the East Coast (see Figure 1; Fowler et al., 2021). Although Western Victoria and the NSW component of the East Coast are considered sustainable, and Eastern Victoria is classified as undefined, the two purely SA snapper stocks are considered depleted due to prolonged recruitment failure and declines in spawning biomass (Fowler et al., 2021). As a result, the Spencer Gulf/West Coast and Gulf St Vincent stocks have been closed to fishing since late 2019. The five southeastern snapper stocks are thought to be mostly sustained by the large spawning aggregations occurring in Spencer Gulf, Gulf St Vincent and Port Phillip Bay (Coutin et al., 2003; Fowler, 2016). However, spawning also occurs in open shelf waters of eastern Vic (as well as further north in NSW), with inlets such as Corner Inlet (located directly eastward of Wilson’s Promontory), Gippsland Lakes (located at Lakes Entrance) and Mallacoota Inlet (located just south of the Vic-NSW border; Neira et al., 2000; Hamer and Jenkins, 2004; Hamer et al., 2005) serving as nurseries.

The spatial boundaries of the five currently recognized snapper stocks in southeastern Australia are based on knowledge of population structure and connectivity gained primarily from otolith microchemistry (Fowler et al., 2005; Hamer et al., 2005; Hamer et al., 2011; Fowler et al., 2017), mark-recapture (Sanders, 1974; Jones, 1984; Coutin et al., 2003; McGlennon, 2003) and demographic analyses (Fowler, 2016). The most detailed population genetic work done in the region are a mitochondrial DNA study with a Victoria component including Port Phillip Bay only (see Figure 1; Donnellan and McGlennon, 1996) and a microsatellite DNA study with Port Phillip Bay as its westernmost sample (Morgan et al., 2018). Additionally, all investigations of population structure and connectivity in snapper in southeastern Australia have focused on one state or sub-region, or involved sparse sampling. Thus far, no population genomic analyses have been done on southeastern Australian snapper. Considering this, and the sustainability issues in SA with regard to resource management, snapper management in the southeast of Australia would benefit from an analysis of genetic structure and connectivity based on comprehensive sampling and genome-wide data.

The marine environment in southeastern Australia is highly complex (see Figure 1). The region possesses large islands, highly variable continental shelf morphology and bathymetry, a topographically complex coastline, temperature fronts, upwellings, boundary currents at the shelf-edge and seasonal inshore wind-driven currents (James and Bone, 2010). It also encompasses three distinct biogeographic provinces (see Figure 1); Flindersia (warm-temperate), Maugea (cold-temperate) and Peronia (warm temperate; Bennett and Pope, 1953). These bioregions coincide with species distributions as well as phylogenetic and population genetic structure (see the reviews: Colgan, 2015; Teske et al., 2017), particularly in taxa without a highly active adult stage such as seaweeds, gastropods, bivalves and echinoderms (e.g., Ayre et al., 1991; Waters et al., 2004; Dawson, 2005; Li et al., 2013; Aguilar et al., 2015; Sinclair et al., 2016; Teske et al., 2017; Muangmai et al., 2022). Consequently, it is not clear how important these marine biogeographic boundaries are for marine fishes that have high dispersal potential across multiple life stages.

Here, we assess population structure in snapper across 11 locations in southeastern Australia using a dataset of genome-wide SNPs. We tested the hypothesis that the population genomic structure of this highly mobile species corresponds with the major biogeographic provinces of the region. We also investigate whether the current spatial scales of assessment and management adopted in the region are consistent with evidence from population genomics. Our dataset includes fish from all currently recognized snapper stocks in southeastern Australia for assessing the presence of genetic breaks at management boundaries (see Figure 1). Lastly, our dataset comprises two temporally spaced samples from two major spawning sites in southeastern Australia for assessing temporal stability in population genomic structure. To our knowledge, our study is the first to use a population genomic framework to assess the importance of the southeastern Australian biogeographic provinces in shaping population structure of a highly mobile marine teleost. It has important implications for the management of some of Australia’s most important snapper fisheries as well as for understanding drivers of population differentiation in coastal marine environments.

## Materials and methods

### Sampling

In 2018 and 2019 a total of 435 adult snapper were sampled from 11 sites along the southeastern Australian coast between Ceduna, South Australia (SA) and Eden, New South Wales (NSW; Figure 1, Table 1). These 11 sites cover all the management areas currently used for snapper in southeastern Australia as well as the vast majority of fishing activity occurring in the region (Fowler et al., 2021), including hotspots near human population centers – the SA gulfs (i.e., Gulf St Vincent and Spencer Gulf) and Victorian embayments (Port Phillip Bay and Western Port Bay). Either muscle or fin samples were taken from the snapper (sampled by authors, volunteers or fisheries researchers at the South Australian Research and Development Institute and the NSW Department of Primary Industries), which were landed by commercial or recreational fishermen (see Table S1). Muscle samples were also obtained from adult snapper used in a past population genetics study that were landed in Northern Gulf St Vincent (*N* = 26) and Port Phillip Bay (*N* = 30) in 2010 and 2011 respectively (Table 1). In our study, these samples were used to assess temporal stability in population structure. Where possible, biological data including age, length and sex were obtained for each sampled individual (Table 1). All samples were preserved in 100% ethanol and stored at −20°C.

**Table 1.** Numbers of individuals (numbers after removing individuals with >20% missing data in parentheses), biological data (average fork lengths in millimeters and ages in years followed by standard deviations in parentheses) and diversity statistics (expected heterozygosity, *H*_E_; observed heterozygosity, *H*_O_; percent polymorphic loci, %PL) for each snapper sample (including the two temporal samples at NGSV and PPB). Refer to Figure 1 for sample provenance (jurisdiction and management area).

|  | **N** | **Avg. FL** | **Avg. age** | **$H_E$** | **$H_O$** | **%PL** |
| --- | --- | --- | --- | --- | --- | --- |
| **Ceduna (CED)** | 37 (37) | 634 (127) | 8.7 (4.1) | 0.187 | 0.188 | 95.0 |
| **Northern Spencer Gulf (NSG)** | 40 (39) | 477 (134) | 3.5 (1.0) | 0.187 | 0.185 | 95.3 |
| **Southern Spencer Gulf (SSG)** | 40 (40) | 444 (149) | 9.1 (4.7) | 0.187 | 0.188 | 95.5 |
| **Northern Gulf St Vincent (NGSV)** | 40 (39) | 488 (184) | 7.0 (3.8) | 0.191 | 0.203 | 95.3 |
| **Northern Gulf St Vincent 2010 (NGSV10)** | 26 (26) | 653 (75) | 9.5 (1.7) | 0.186 | 0.185 | 91.8 |
| **Southern Gulf St Vincent (SGSV)** | 40 (40) | 665 (139) | 10.3 (3.5) | 0.189 | 0.187 | 96.7 |
| **Kingston SE (KSE)** | 38 (38) | 514 (98) | 7.9 (2.5) | 0.188 | 0.185 | 94.8 |
| **Portland (PLD)** | 40 (39) | 331 (42) | N/A | 0.187 | 0.184 | 93.6 |
| **Port Phillip Bay (PPB)** | 40 (40) | 592 (61) | N/A | 0.189 | 0.189 | 94.4 |
| **Port Phillip Bay 2011 (PPB11)** | 30 (30) | 504 (69) | 8.9 (1.3) | 0.187 | 0.186 | 91.5 |
| **Western Port Bay (WPB)** | 40 (40) | 438 (139) | 6.5 (3.3) | 0.188 | 0.191 | 94.4 |
| **Lakes Entrance (LE)** | 40 (40) | 337 (77) | N/A | 0.187 | 0.187 | 95.6 |
| **Eden (EDN)** | 40 (40) | 283 (31) | 2.7 (1.5) | 0.183 | 0.181 | 95.4 |

### DNA extraction, genomic library preparation and sequencing

DNA was extracted from each tissue sample using a modified salting out protocol (Sunnucks and Hales, 1996). Each DNA sample was subsequently quantified with a Qubit v2.0 (Life Technologies) and diluted to 20 ng/μL. Diluted DNA was then checked for quality with agarose gel electrophoresis on a 1% TBE gel. Library preparation for double digest restriction-site associated DNA (ddRAD) sequencing was carried out as in Peterson et al. (2012), with modifications as detailed in (Brauer et al., 2016). Briefly, ~200ng of DNA per sample was digested using the restriction enzymes Msel and Sbfl-HF (New England Biolabs). Subsequently, ligation of each digested DNA sample to one of 96 unique 6 bp barcodes was performed before pooling into groups of 12 samples. Pippin Prep (Sage Science) was then used to select DNA fragments within each pool between 300 and 800 bp. PCR was then employed to amplify the resulting DNA fragments in each pool, which were separated into three 25 μL reactions to minimise PCR artefact bias. PCR products for each pool were then combined and their size distributions assessed using a 2,100 Bioanalyser (Aligent Technologies). Pools were then quantified with Qubit and aliquots of equal concentrations subsequently combined in equal concentrations, resulting in DNA libraries of 96 samples. Sequencing was done on six lanes of an Illumina Hi-seq 4,000 (150 bp paired end) at Novogene (Hong Kong). Each 96-sample library included six replicates for assessing sequencing and genotyping errors.

### Bioinformatics and characterisation of putatively neutral SNPs

Raw sequence reads were put through a bioinformatic pipeline as in Bertram et al. (2022) to produce a high-quality SNP dataset. Briefly, raw data was demultiplexed using the *process_radtags* module of STACKS 2.0 (Catchen et al., 2013). Subsequently, restriction sites, barcodes and RAD tags were trimmed from sequences with TRIMMOMATIC 0.36 (Bolger et al., 2014). The resulting sequence reads were aligned to the snapper genome of Catanach et al. (2019) with BOWTIE 2 (Langmead and Salzberg, 2012). SNPs were then called using BCFTOOLS 1.16 (Narasimhan et al., 2016). The resulting SNP dataset was then filtered using VCFTOOLS 0.1.16 (Danecek et al., 2011). Specifically, initial filtering was conducted to retain just bi-allelic SNP markers occurring in >80% of individuals in all populations and with a minimum minor allele frequency of 0.03, as well as to remove indels. Individuals with >20% missing data after completing these filtering steps were removed from the original unfiltered SNP dataset and the initial filtering steps were conducted again on the dataset with fewer individuals (i.e., those with <20% missing data). Subsequently, several other filtering steps were performed to remove SNPs with low read and mapping quality, SNPs with very high coverage, SNPs departing from Hardy-Weinberg equilibrium in >2/3 of locations, SNPs with very high call error rate and physically linked SNPs (see Table S2).

Lastly, SNP markers potentially under directional selection were identified and removed from the dataset to leave a putatively neutral one. Outlier SNPs were identified using the Bayesian method of BAYESCAN 3.0 (Foll and Gaggiotti, 2008), which was carried out initially with 20 pilot runs (with 5,000 iterations for each), followed by 100,000 iterations including a burn-in period of 50,000 iterations. A false discovery rate of 5% with a prior odd of 10 was employed to identify outlier SNPs, which were subsequently removed from the dataset. A separate study on snapper is utilizing candidate adaptive SNPs to investigate adaptation associated with environmental variation (Brauer et al. unpublished).

### Genomic diversity statistics and pairwise genetic differentiation (*F*_ST_)

Levels of genomic diversity in each snapper sample was assessed with estimates of expected and observed heterozygosity (*H*_E_ and *H*_O_, respectively) and percent polymorphic loci (%PL) using the *populations* module in STACKS 2 (Rochette et al., 2019). Estimates of genetic differentiation (*F*_ST_) between pairs of samples were calculated with ARLEQUIN 3.5 (Excoffier and Lischer, 2010), using 1,000 permutations to assess significance. *P*-values were subsequently corrected for multiple comparisons with the false discovery rate method using the R package BASE 4.0.3 (Team, 2021). Global *F*_ST_ was also determined using HIERFSTAT 0.5-10 (Goudet et al., 2015) in R, with 95% confidence intervals (CIs) calculated using 1,000 permutations.

### Clustering analyses

Principal components analysis (PCA) was performed using VEGAN 2.5-6 (Oksanen et al., 2018) in R. Missing genotypes (0.9% of data matrix) were replaced with the most common one at the locus. Next, the number of genetically distinct groups was inferred using the maximum likelihood approach of ADMIXTURE 1.3 (Alexander et al., 2009; Alexander and Lange, 2011). The software’s cross-validation procedure was used to determine the most likely number of genetic clusters in the dataset (i.e., *K*). A 5-fold cross-validation was conducted for values of *K* between one and eight. Membership probabilities were visualized graphically using GGPLOT2 3.3.3 (Wickham, 2016) in R. The analysis was also run with the two temporal samples to investigate temporal variation in membership probabilities to each inferred genetic cluster.

### Isolation by coastal distance

The relationship between the genetic differentiation between samples (linearized *F*_ST_, i.e., *F*_ST_/(1-*F*_ST_)) and coastal distance (i.e., isolation by distance; IBD) was investigated using Mantel tests in GENALEX 6.5 (Peakall and Smouse, 2012). Coastal distances were estimated as the shortest distance between sampling locations following the coastline using the *viamaris* function in MELFUR 0.9 (https://github.com/pygmyperch/melfuR). Mantel tests were conducted with all samples as well as separately for each of the two broad genetic groupings identified with the clustering analyses (i.e., the SA and Vic groups). Significance of the three Mantel tests was determined with 10,000 permutations.

### Spatial autocorrelation

The spatial autocorrelation of samples was investigated first with spatial principal component analysis (sPCA) using ADEGENET 2.1.5 in R (Jombart, 2008). This method is optimized to reveal cryptic spatial patterns of genetic differentiation and produces principal component scores summarizing both the non-spatial variability and the spatial autocorrelation (calculated as Moran’s *I*; Moran, 1948; Moran, 1950) among genotypes. For this analysis, we used a connection network based on a spatial weights matrix generated from the inverse of the absolute coastal distances between sampling locations. The analysis was run on the whole 2018/19 dataset and then separately for the two large groups identified by the clustering analyses (i.e., the SA and Vic groups). The presence of positive spatial autocorrelation in the three different datasets was assessed using 9,999 permutations. Lagged spatial principal component scores were plotted for improved clarity of spatial patterns using the R package ADE4 1.7-18 (Dray and Dufour, 2007).

Second, spatial autocorrelation coefficients (*r*) were calculated for each site separately using GENALEX 6.5 (Smouse and Peakall, 1999; Peakall and Smouse, 2012) to investigate within-location spatial autocorrelation. The significance of *r* values was calculated using 1,000 bootstraps and 95% CIs around the null hypothesis of randomly distributed genotypes were calculated using 1,000 permutations. To avoid overinterpretation of spatial autocorrelation coefficients, we regarded a value of *r* to be significant only if it occurred outside of the 95% CIs around the null hypothesis of zero correlation and if its error bars did not overlap the x-axis.

## Results

### SNP genotyping and genomic diversity

Following alignment of sequence reads to the snapper genome and SNP calling, ~7.3 million raw variants were discovered. Three individuals were removed from the dataset due to having >20% missing data, leaving 488 for subsequent analyses (432 from 2018/19 and 56 from 2010/11; Table 1). After completing all quality filtering steps and removing 350 SNPs putatively under selection, 10,916 SNPs remained (details in Table S2). Average missing data per sample was 1.0% (range: 0-13.1%) and average coverage depth per locus per sample was 102.6 (range: 6.9-303.5).

Genetic diversity was highly similar across the 11 snapper localities collected in 2018 and 2019 (see Table 1), with expected heterozygosity (*H*_E_) ranging between 0.183 (EDN) and 0.191 (NGSV), observed heterozygosity (*H*_O_) ranging between 0.181 (EDN) and 0.203 (NGSV) and percent polymorphic loci (%PL) ranging between 93.6% (PLD) to 96.7% (SGSV).

### Clustering analyses

Although *K* = 2 was the most supported by ADMIXTURE (Figure S1), *K* = 3 closely followed (Figure 2), and the PCA indicated the presence of three clusters (Figure S2). These three groups, herein referred to as the South Australia (CED to SGSV; SA), Victoria (KSE to LE; Vic) and New South Wales (EDN; NSW) stocks, correspond with the three biogeographic provinces in southeastern Australia – Flindersia, Maugea and Peronia (Figures 1 and 2). In the SA stock, two Vic migrants and several individuals with mixed ancestry occurred, but only in Gulf St Vincent (NGSV and SGSV), adjacent to the stock’s eastern boundary. In the other direction, three SA migrants were detected in the Vic stock within the sample adjacent to the stock’s western boundary (i.e., KSE). While an abrupt genetic break occurred between the SA and Vic stocks, considerably more overlap occurred between the Vic and NSW stocks, suggesting that a region of isolation by distance (IBD) may occur between them (Figure 2). Even though LE clearly clustered with the Vic stock, our sample contained both NSW migrants and a considerable proportion of individuals with mixed ancestry. WPB was the westernmost Vic sample containing individuals with considerable NSW ancestry. Although the EDN sample contained predominantly individuals with NSW ancestry, it also contained fish that were migrants from Vic and that had mixed ancestry.

**Figure 2.**
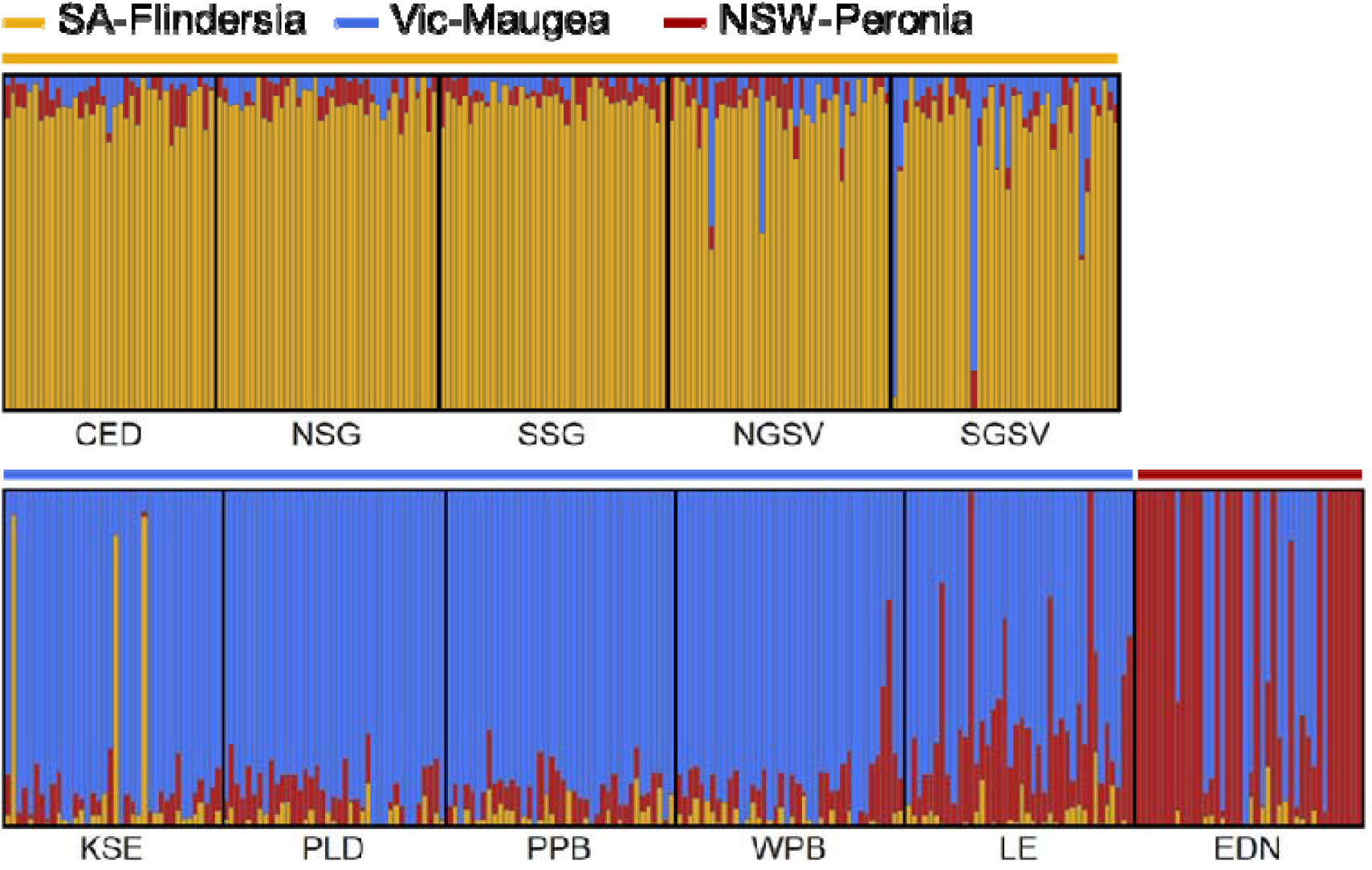
ADMIXTURE results for *K* = 3 based on the 432 snapper individuals (excluding the temporal samples from NGSV and PPB) and 10,916 neutral SNPs. Biogeographic provinces (Flindersia, Maugea, Peronia) and genetic groupings (SA, Vic, NSW groups) are marked above the plots. Each vertical bar represents an individual and its probability of membership to each of the *K* groups is indicated by its colour makeup.

### Temporal analysis

Comparisons of ADMIXTURE ancestry proportions across the temporally spaced samples at NGSV and PPB indicated stability in population genetic structure over periods of approximately nine and eight years, respectively (Figure 3). Additionally, the analysis suggested that dispersal from the Vic stock into Gulf St Vincent is recurrent since one Vic migrant was detected by ADMIXTURE in the NGSV10 sample (see Vic stock outlier in NGSV10 sample in Figure 3).

**Figure 3.**
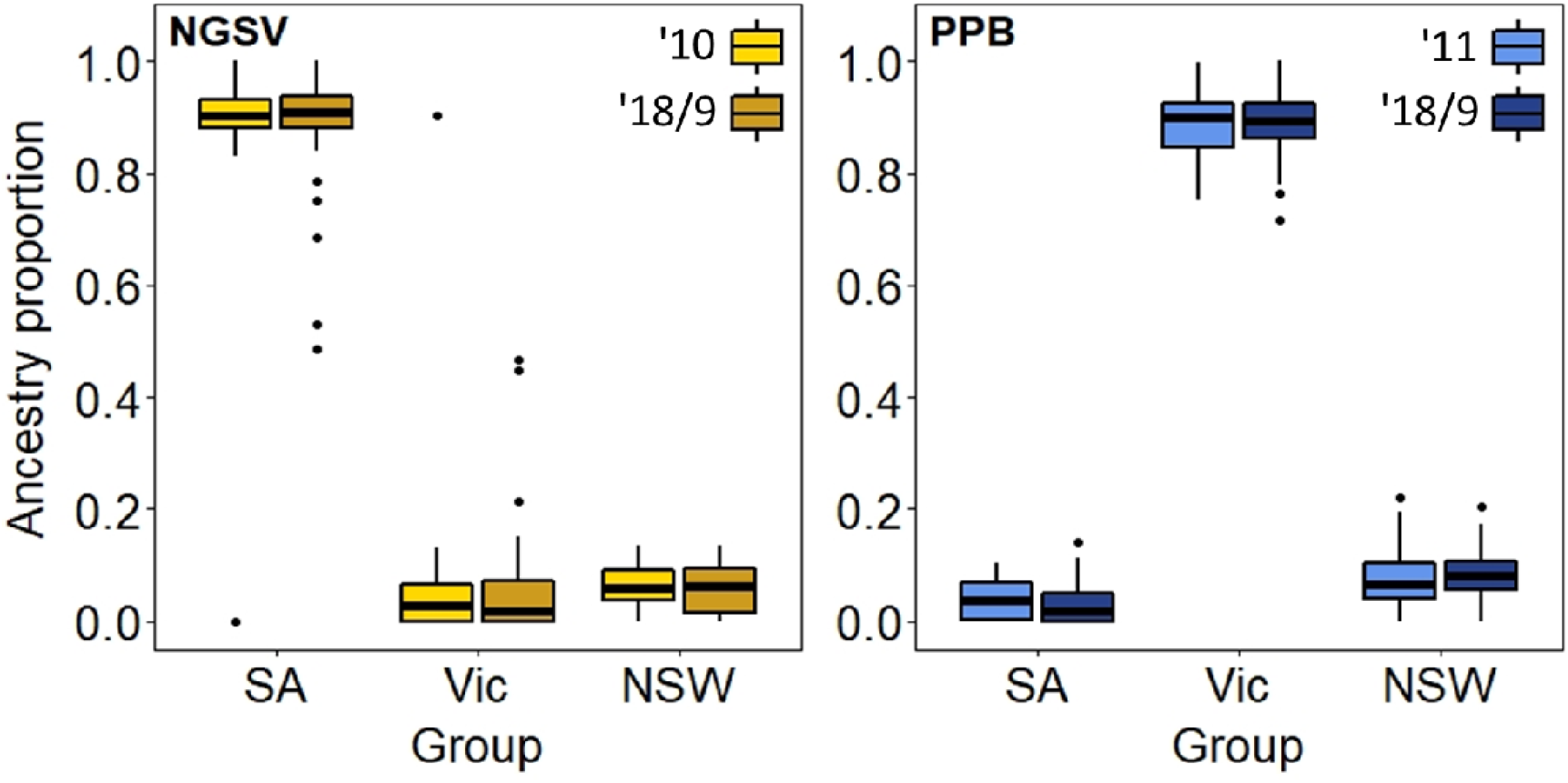
Temporal comparisons of snapper membership to the three clusters identified by ADMIXTURE between the two sampling periods at Northern Gulf St Vincent (NGSV; gold) and Port Phillip Bay (PPB; blue). None of the comparisons at either location were significantly different, suggestive of stability in patterns of population genetic structure.

### Pairwise *F*_ST_ and isolation by coastal distance (IBD)

Global *F*_ST_ was low at 0.0116 (95% CIs: 0.0098, 0.0135), and pairwise *F*_ST_ comparisons were also low, ranging between 0.0002 (CED:SSG) and 0.0223 (NSG:PLD; Figure 4). Out of the 55 pairwise *F*_ST_ values, 42 were significant after correction for multiple comparisons through the FDR method. The two previously identified broad groupings (i.e., SA and Vic stocks) were clearly discernible in the pairwise *F*_ST_ heatmap, as well as the differentiation of EDN. NGSV and LE were the only samples to show significant differentiation within their respective clusters.

**Figure 4.**
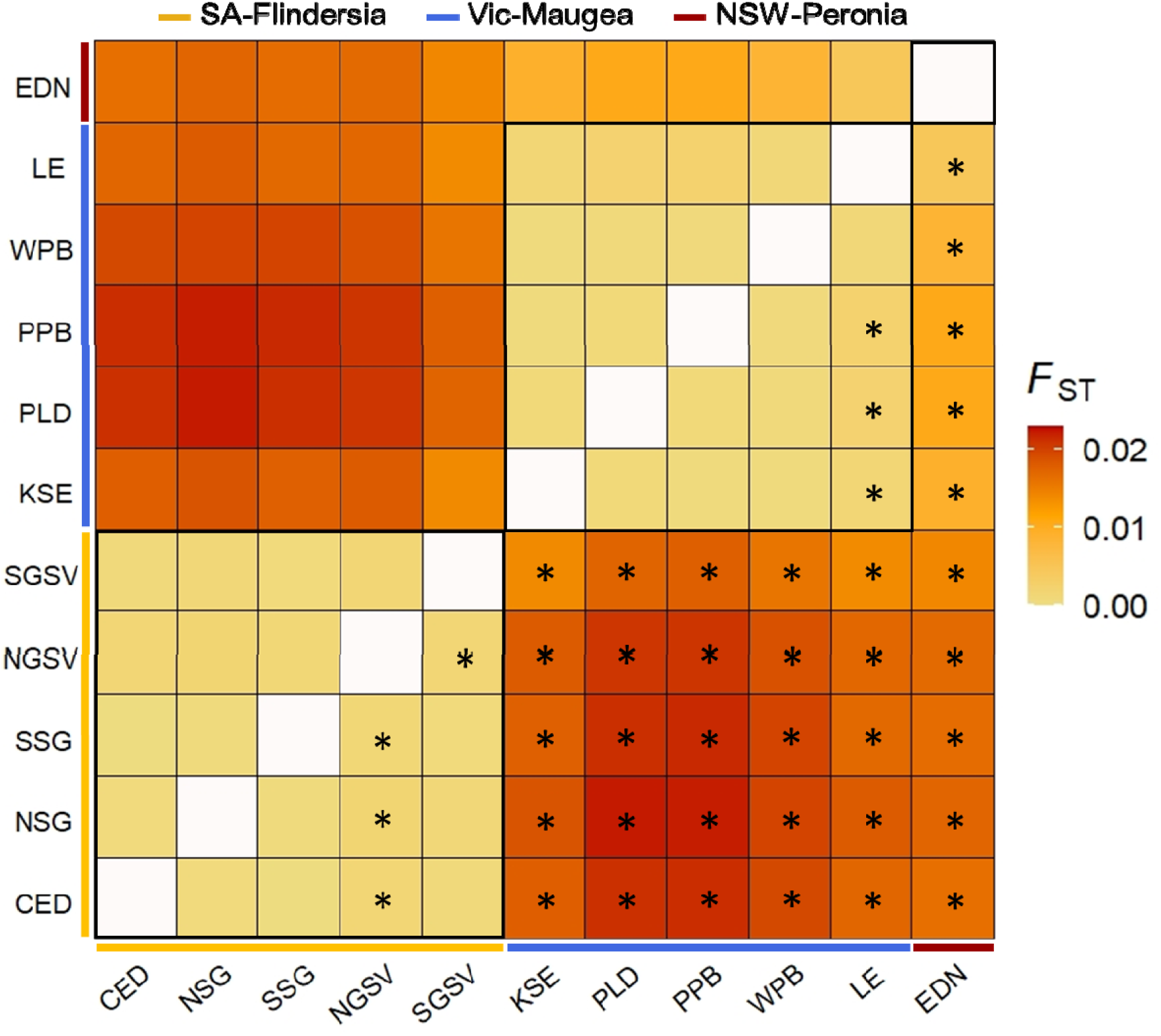
Heatmap of pairwise *F*_ST_ values (i.e., genetic differentiation) between all snapper samples (except the two temporal samples) based on the 10,916 neutral SNPs, with significant comparisons denoted with an asterisk in the lower diagonal (42 out of 55 comparisons). Main groupings (i.e., CED to SGSV; KSE to LE; EDN) and biogeographic provinces (Flindersia, Maugea, Peronia) are marked with black squares and coloured lines. *F*_ST_ values ranged between 0.0002 and 0.0223, and global *F*_ST_ was 0.0116.

A Mantel test uncovered significant IBD between the west coast of SA and the south coast of NSW (r = 0.57, *p*-value = 0.003; Figure S3). However, this trend was largely driven by differentiation at genetic stock boundaries since no IBD occurred within the SA and Vic stocks (Figure S3).

### Spatial autocorrelation

Significant global spatial structures were detected in the whole dataset (*p*-value = 0.0001) as well as when the SA and Vic stocks were considered separately (*p*-values = 0.0109 and 0.0265, respectively). The sPCA including all 2018/9 samples differentiated the three groups identified with ADMIXTURE and PCA (i.e., the SA, Vic and NSW stocks; Figure S4). Additionally, they illustrated the nature of the boundaries between the groups, with the SA-Vic boundary occurring abruptly between SGSV and KSE and a transition region (i.e., a region of IBD) occurring from WPB to EDN between the Vic and NSW stocks (concordant with the ADMIXTURE results). The sPCA of the SA stock differentiated Gulf St Vincent (and especially NGSV) from all other SA sites (i.e., NSG, SSG and CED; Figure 5a) and the sPCA of the Vic group differentiated LE from all other Vic sites (i.e., WPB, PPB, PLD and KSE; Figure 5b).

**Figure 5.**
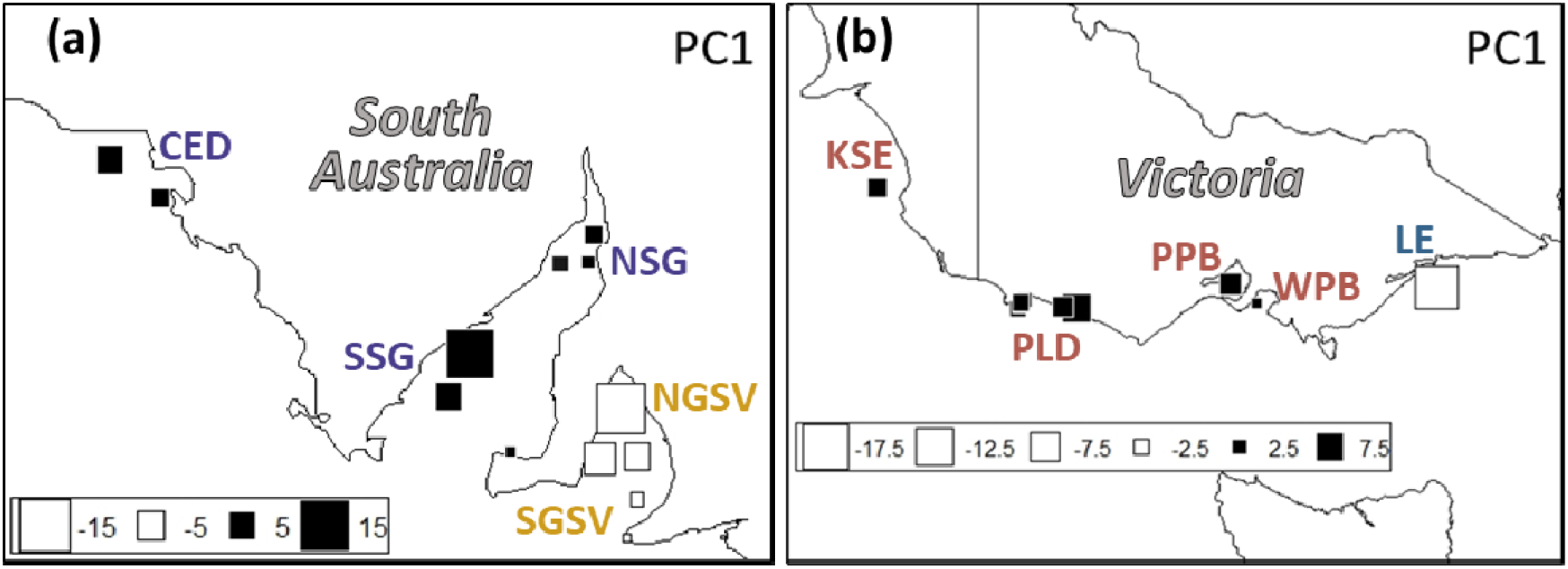
The first global scores of spatial PCA of the (a) South Australian snapper group and the (b) Victorian snapper group, which were both based on 10,916 neutral SNPs. Site names are coloured according to the management area in which they reside (as in Figure 1): 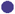 = Spencer Gulf/West Coast,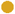 = Gulf St Vincent, 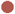 = Western Victoria, 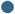 = Eastern Victoria, 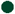 = East Coast.

Within site, spatial autocorrelation indicated that genotypes were significantly correlated, suggesting that considerable local recruitment and/or site fidelity occurs at all sites except SSG, PLD and WPB. Spatial autocorrelation coefficients were highest for NSG, NGSV, LE and EDN, and the most variable for NSG, NGSV, PLD, PPB and EDN (Figure 6).

**Figure 6.**
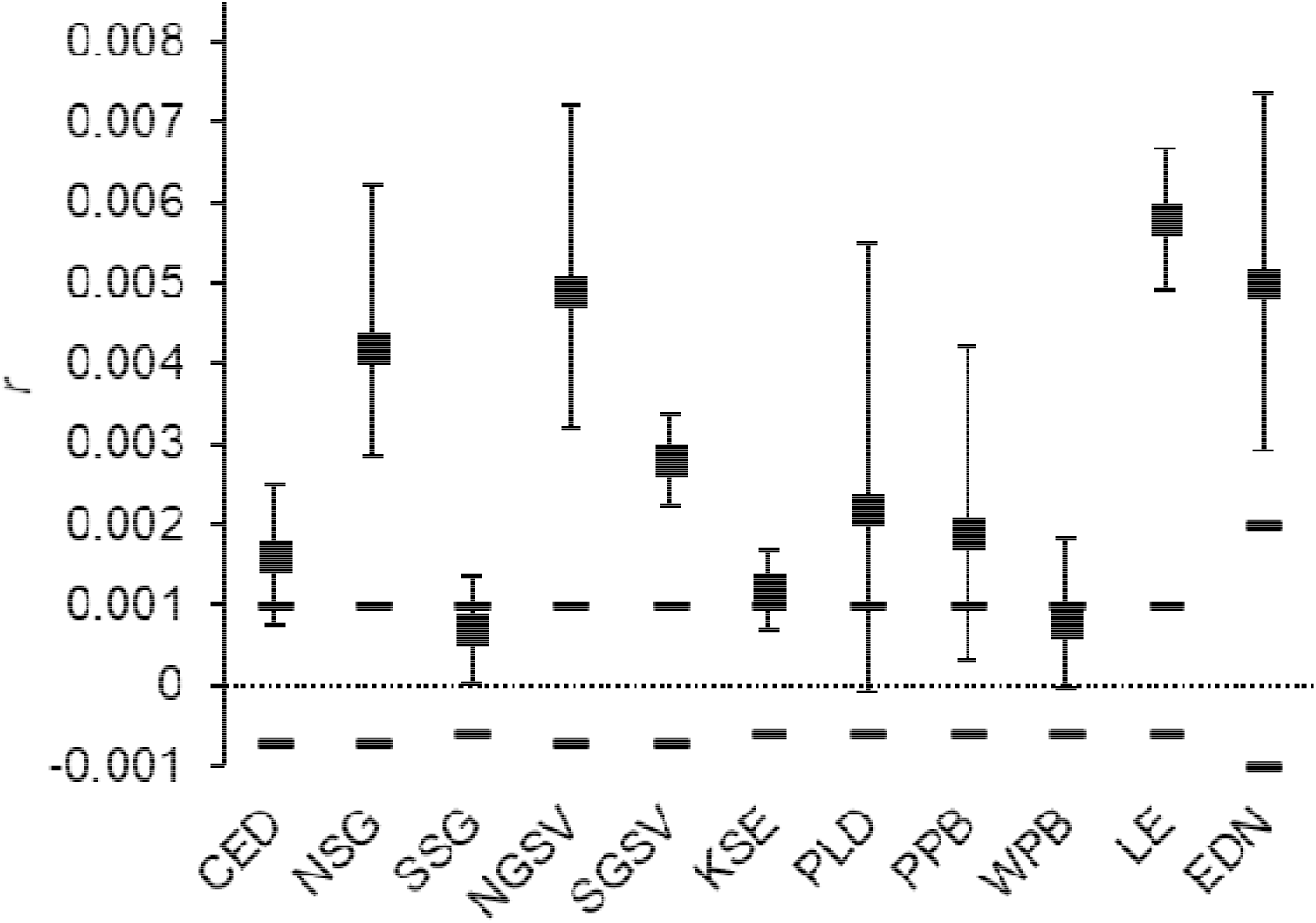
Within site genotypic autocorrelation coefficients (*r*) for snapper in southeastern Australia with 95% confidence intervals. Individuals are more similar than expected by chance at all sites except for at SSG, PLD and WPB.

## Discussion

Using comprehensive sampling and a genome-wide SNP dataset, we uncovered genetic breaks in the heavily exploited teleost snapper across southeastern Australia concordant with the region’s biogeographic barriers. Although these genetic breaks coincide with only two of the four management boundaries used for snapper in the region, we found some evidence for fine-scale genetic structure corresponding with the two other boundaries. This fine-scale structure likely reflects spatial variation in spawning and recruitment dynamics and highlights the power of genome-wide SNP datasets for revealing evolutionary and demographic processes in highly abundant and dispersive species. Our study also confirms the importance of marine biogeography in shaping intraspecific genetic structure, even in species with high dispersal potential like snapper.

### Bioregional divisions and stock structure

Coastal marine species often exhibit distinct breaks within their distributions where movement is limited, frequently aligning with bioregional boundaries (Bennett and Pope, 1953; Briggs, 1974; Kelly and Palumbi, 2010; Sivasundar and Palumbi, 2010; Colgan, 2015; Teske et al., 2017). These boundaries usually coincide with shifts in features like ocean circulation, bathymetry, temperature, coastline topography and habitat (Hayden and Dolan, 1976; Gaylord and Gaines, 2000; Sivasundar and Palumbi, 2010; Waters et al., 2010; Barton et al., 2018). At a broad scale, the population structure of snapper in southeastern Australia is predominately shaped by two genetic discontinuities (i.e., the Murray River Mouth and the Vic-NSW border) which correspond to the boundaries between the three biogeographic provinces in the region – Flindersia, Maugea and Peronia (see Figure 1). Coastal distance was not a dominant driver of genetic distance for snapper in this region, which contrasts with recent work done in the western part of the species’ range (Bertram et al., 2022). Very few published population genetic studies on finfish in southeastern Australian with high dispersal potential across all life stages have reported structuring corresponding to these biogeographic boundaries. Population genetic work (employing mitochondrial or microsatellite DNA markers) with mulloway (*Argyrosomus japonicus*), gemfish (*Rexea solandri*) and round herring (*Etrumeus sadina*) have reported structuring broadly coinciding with the boundary between the Maugea and Peronia biogeographic provinces (Colgan and Paxton, 1997; DiBattista et al., 2014; Barnes et al., 2015). To our knowledge, the only other finfish shown to exhibit a genetic discontinuity at the boundary between Flindersia and Maugea is the black bream (Burridge and Versace, 2007). However, this sparid is expected to have low dispersal potential relative to a species like snapper since it can complete its entire life cycle in estuaries and only occurs in coastal marine environments during periods of high river discharge or anoxia (Lenanton, 1977; Holt, 1978; Sherwood, 1982; Lenanton, 1999).

The relatively strong genetic break detected in the vicinity of the Murray River Mouth in southeast SA corresponds with the transition zone between the warm temperate Flindersia province and the cold temperate Maugea province. This finding is concordant with previous work on snapper involving otolith microchemistry (Fowler et al., 2017), demographic analyses (Fowler, 2016), mark-recapture (Sanders, 1974; Coutin et al., 2003), acoustic telemetry (Lédée et al., 2021), allozyme and mitochondrial DNA markers (MacDonald, 1980; Donnellan and McGlennon, 1996) and microsatellites (Gardner et al., 2022). The most detailed population structure work done in the region used otolith microchemical analyses and concluded that the southeast of SA (east of Kangaroo Island) is dominated by fish resulting from spawning events in Port Phillip Bay (Fowler et al., 2017). Additionally, Fowler et al. (2017) indicated that the Gulf St Vincent and Spencer Gulf/West Coast stocks are dominated by fish resulting from spawning events in the northern gulfs. Whilst otolith microchemistry work (Fowler et al., 2017) and demographic analyses (Fowler, 2016) indicated that dispersal occurs from the Vic stock to just south of of Gulf St Vincent, we were also able to detect dispersal into the gulf. Demographic analyses suggest that such broad-scale dispersal results from very strong recruitment events in Port Phillip Bay (Fowler, 2016).

In contrast to the sharp genetic break detected near the Murray River Mouth, a region of admixture occurred at the transition between the cold temperate Maugea province and the warm temperate Peronia province. This region, between Wilson’s Promontory and the south coast of NSW, is a heterogeneous mix between Vic and NSW snapper. Although fewer studies on snapper have included the lower east coast, the inferred admixture was also found in a microsatellite analysis focusing on eastern Australia (Morgan et al., 2018). Additionally, otolith microchemistry analyses and recruitment surveys suggest that eastern Victorian snapper (east of Wilson’s Promontory to the NSW border) originate from a number of different sources including local inshore areas (e.g., off Lakes Entrance), Port Phillip Bay and areas further north along the east coast (Hamer and Jenkins, 2004; Hamer et al., 2005; Hamer et al., 2011). Our results indicate that dispersal is more bi-directional (as well as more frequent) between the Vic and NSW stocks than between the SA and Vic stocks. This implies that considerable numbers of Vic snapper make north-eastward movements, opposite to the direction of the East Australian Current (EAC). Other marine species, including the common dolphin (*Delphinus delphis*), estuary perch (*Macquaria colonorum*), round herring (*Etrumeus teres*), sardine (*Sardinops sagax*), eastern king prawn (*Penaeus plebejus*) and tailor (*Pomatomus saltatrix*), are also known to make northward movements along the east coast of Australia (Montgomery, 1990; Ward et al., 2003; Möller et al., 2011; Shaddick et al., 2011). Such dispersal events are thought to occur due to northward flowing currents generated by EAC eddies (Ridgway and Dunn, 2003; Oke et al., 2019) and/or patterns of adult migration. Indeed, tagging data indicate that some sub-adult and adult snapper make northward movements along the east coast of Australia (Sanders, 1974; Coutin et al., 2003; Stewart et al., 2019).

Differentiated populations corresponding with the biogeographic provinces may represent groups that were isolated to different refugia during glacial maxima when continental shelf habitat was dramatically reduced as well as fragmented (Waters and Roy, 2003; Waters, 2008; James and Bone, 2010; Sinclair et al., 2016; Dolby et al., 2018). When sea levels were ~129 m below their present level during the last glacial maximum (LGM), the Bassian Isthmus (between Tasmania and mainland Australia) and Lacepede Shelf (east of Kangaroo Island in the southeast of SA) were exposed, thereby acting as gene flow barriers between the provinces. Why genetic differentiation has persisted since the inundation of these regions is uncertain. Most population genetic studies done in the region have involved reef associated organisms like intertidal invertebrates and macroalgae (see Colgan, 2015; Teske et al., 2017). As a result, many suggest that the genetic discontinuities occurring at the bioregion boundaries are maintained due to areas of unsuitable sandy habitat at the Coorong (located between Kingston SE and the Murray River Mouth) and Ninety Mile Beach (spanning from just east of Wilson’s Promontory to around Lakes Entrance). However, snapper inhabit both sandy and reef habitats and certainly possess the physical ability to migrate between these broad areas. Other studies on species associated with sandy areas, including seagrass (*Posidonia australis*), an asteriid sea star (*Coscinasterias muricata*) and cuttlefish (*Sepia apama*), have found intraspecific population structure coinciding with one or more of the bioregion boundaries (e.g., Kassahn et al., 2003; Waters and Roy, 2003; Sinclair et al., 2016). Alternatively, the maintenance of such historic differentiation could be due to complex flow patterns on the lower east coast (in the case of the Maugea-Peronia boundary; i.e., the convergence of the Leeuwin Current and EAC, and EAC eddies) as well as competitive or density-dependent interactions (Hidas et al., 2007; Ayre et al., 2009; Waters, 2011; Waters et al., 2013; Teske et al., 2017). The latter is perhaps particularly likely since snapper are highly fecund and abundant. Large amounts of immigration would therefore be required to cause genetic changes in a large local population, which may be hindered by local resource monopolization and competitive superiority of locally adapted fish (De Meester et al., 2002; Fraser et al., 2015).

### Fine-scale structure within stocks

Spatial autocorrelation analyses can identify the spatial scales over which individuals are more genetically similar than expected at random and therefore can indicate whether recruitment occurs predominately locally and/or whether site fidelity occurs post settlement. Our spatial autocorrelation analyses indicate that genomic variation was non-randomly distributed within both the SA and Vic stocks. In SA, sPCA showed subpopulation-level structure between the Gulf St Vincent (especially the north of Gulf St Vincent) and the Spencer Gulf and Ceduna. Additionally, the site-specific spatial autocorrelation analysis suggested that within the SA stock, individuals were the most genetically similar at Northern Spencer Gulf and Northern Gulf St Vincent. These two locations are thought to be the most important spawning and nursery areas for snapper in SA (Fowler and Jennings, 2003; Fowler et al., 2005). During the summer when snapper spawn in SA (Saunders et al., 2012), temperature fronts at the entrances of the gulfs largely prevent exchanges with the shelf (Bruce and Short, 1990; Vaz et al., 1990; Petrusevics, 1993; Teske et al., 2015). As a result, larvae resulting from spawning in the gulfs are largely expected to settle locally, consistent with our spatial autocorrelation results. In addition, the spatial autocorrelation results may reflect demographic differences between the two gulfs reported in analyses of age composition and recruitment history (Fowler, 2016), as well as the limited movement of adults found by mark-recapture work (Jones, 1984; McGlennon, 2003). These analyses suggest that juveniles resulting from periodic strong recruitment events in either of the two gulfs dominate catches across the entire geographical region corresponding with the SA stock after recruiting to the fishery at ~3 years of age. However, when these infrequent events do not occur, the recruitment dynamics of the two gulfs appear largely independent (Fowler, 2016). Additionally, the majority of fish tagged in Spencer Gulf and Gulf St Vincent have been recaptured within 37km of their release sites, with only a couple of individuals observed to move between the two gulfs (SGSV to NSG; Jones, 1984; McGlennon, 2003). As a result, adult site fidelity may be contributing to the spatial autocorrelation results. Overall, these results exemplify the power of population genomics to detect signal consistent with distinct spawning aggregations and local recruitment in abundant and highly dispersive fisheries resources.

Despite Spencer Gulf and Gulf St Vincent supporting different spawning aggregations (Fowler, 2016), which reproduce simultaneously during the summertime, as well as the small differences inferred with sPCA, the two localities form part of the same genetic stock. This contrasts with Western Australia and Vic, where major spawning sites are represented by separate genetic stocks (this study and Bertram et al., 2022). It is possible that the Spencer Gulf and Gulf St Vincent spawning groups originated from a single ancestral coastal population during the LGM when the gulfs were dry (James and Bone, 2010). Colonization following the inundation of the gulfs ~10,000 years ago likely then allowed for the formation of spawning groups observed today. The intermittent demographic isolation presently occurring over short time scales between the gulfs is likely not prolonged enough to cause the degree of population genetic differentiation uncovered between the SA, Vic and NSW genetic stocks. Additionally, the abundances of snapper in these areas would have to be greatly reduced for a very prolonged period to produce significant genetic differentiation.

Outside of the SA stock, we detected a pattern of positive spatial autocorrelation along the lower east coast between Port Phillip Bay and Eden indicative of isolation by distance. This result is consistent with those obtained with ADMIXTURE and an allozyme study by Meggs et al. (2003). The site-specific spatial autocorrelation analysis indicated that individuals were most similar at the two most eastern locations (i.e., Lakes Entrance and Eden), despite them being quite heterogeneous, containing individuals with a range of ancestry profiles (Figure 2; Figure 6). At Lakes Entrance and Eden, spawning occurs outside of coastal embayments and therefore eggs and larvae are subjected to oceanic circulation patterns. Nevertheless, the positive spatial autocorrelation uncovered at the two sites may relate to local retention of pelagic life stages by EAC eddies (Mullaney and Suthers, 2013). Another possibility is that the abundance of snapper at Lakes Entrance and Eden is relatively low, increasing the chance of sampling genetically alike individuals. This idea is supported by the lower catches of snapper off Lakes Entrance and Eden (however note their distances from human population centers in Figure 1; Stewart, 2020; Fowler et al., 2021). The lack of large embayments (for supporting large spawning aggregations and recruits) as well as the scarcity of rocky reefs (preferred adult habitat), may prevent populations from becoming very large in these areas.

### Temporal analysis supports both broad and fine-scale genetic patterns

The temporal analysis indicated stability in admixture proportions at Northern Gulf St Vincent and Port Phillip Bay over an approximately eight-year period. These results indicate the persistence of the genomic signal of local recruitment and fidelity of adults associated with these two important spawning sites. This result is concordant with similar work done in the western part of the range of snapper at the major spawning site Cockburn Sound (Bertram et al., 2022). Despite the smaller size of the 2010 Northern Gulf St Vincent sample compared to the 2018/19 one (26 vs 39 individuals), it still captured dispersal from the Vic stock into the gulf in the form of a full migrant. This suggests that the dispersal events detected in our 2018/19 samples are temporally persistent (although note that such dispersal probably also depends on recruitment success in the Vic stock, which is highly variable between years; Hamer and Jenkins, 2004). The two temporally spaced Port Phillip Bay samples were both devoid of migrants from other stocks as well as admixed individuals. Our temporal analysis therefore not only suggests temporal stability in the spawning groups and recruitment at these two sites, but also consistency in the patterns of migration between adjacent stocks.

### Management implications

Our detailed assessment of the population genomic structure of snapper in southeastern Australia allows us to assess the correspondence between current spatial scales of assessment and management in the region and both broad and fine-scale genetic structure. Our results provide strong support for the management boundaries already in place at the Murray River mouth (i.e., the boundary between the Gulf St Vincent and Western Victoria stocks) and at the Vic-NSW border (i.e., the boundary between the Eastern Victoria and East Coast stocks), agreeing with previous work utilizing mark-recapture, otolith microchemistry as well as analyses of mitochondrial DNA, allozyme and microsatellite variation. Importantly, population genomics also supports the spatial scales of the SA fishery closure initiated due to spawning biomass depletions in Gulf St Vincent and Spencer Gulf (Fowler et al., 2021). The Spencer Gulf/West Coast and Gulf St Vincent stocks were closed to commercial and recreational fishing in late 2019 for a period of three years. Our results support the inclusion of the west coast of SA and the exclusion of southeast SA in the ban since they indicate that the former is dependent on spawning in the gulfs (particularly Spencer Gulf), while the latter is dependent on spawning in Vic. The 2022 SA snapper stock assessment indicated that the Spencer Gulf/West Coast and Gulf St Vincent stocks are not recovering, while biomass is increasing in southeast SA (Drew et al., 2022). These results further corroborate our conclusion that the productivity of snapper in southeast SA is not supported by spawning events occurring in the SA gulfs.

The management boundaries currently in place between the two SA gulfs and at Wilson’s promontory are not marked by genetic breaks. However, we did detect fine-scale genetic structure corresponding to the two boundaries, likely related to spatial variation in spawning and recruitment dynamics. These management boundaries therefore may be suitable depending on the spatial management objectives. Particularly, the unique spawning and recruitment dynamics occurring in the two SA gulfs and eastern Vic could mean that if depletions occur in these areas, recovery through gene flow from adjacent populations may occur very slowly. Additionally, replenishment would depend on both the abundances of snapper and the frequency of strong recruitment events in adjacent populations, with the latter being highly temporally variable in SA and Vic (Hamer and Jenkins, 2004; Fowler, 2016). Although strong recruitment has occurred recently in the Vic stock (Port Phillip Bay in 2018), recent spawning biomass reductions and prolonged recruitment failure in the SA gulfs could mean that spawning groups in either Spencer Gulf or Gulf St Vincent may not be able to recover the snapper resources within the entire area currently closed to fishing in a timeframe relevant to fisheries management (Fowler, 2016). Since the sampling strategy for this study was conceived with the aim of assessing broad-scale population genetic differentiation, more focused fine-scale sampling and analyses may be beneficial for gaining a more detailed understanding of small-scale genetic processes. Additionally, an analysis of local adaptation (e.g., based on seascape genomics) would contribute to understanding of patterns of gene flow and recruitment within the two genetic stocks.

## Conclusion

Our study highlights that the well-known southeastern Australian biogeographic provinces can match intraspecific patterns of genetic structure in highly dispersive species like snapper. Traditionally, population differentiation coinciding with these biogeographic provinces was thought to occur (or at least be maintained) due to the vast areas of sandy habitat around the Coorong in SA and Ninety-mile beach in Victoria. However, snapper occur in both sandy and reef habitats and therefore it is possible that other forces may be important for shaping intraspecific distribution patterns. Two of the current management boundaries used for snapper in the region coincided with the genetic discontinuities detected at bioregional boundaries. Although the remaining two management boundaries did not coincide with distinct genetic breaks, they were marked by finer-scale genetic structure. Our study highlights the value of population genomic surveys involving exploited marine species with high dispersal potential for uncovering both strong genetic breaks and local-scale structuring associated with spatial variation in spawning and recruitment dynamics.

## Acknowledgements

This research was financially supported by the Australian Research Council (LP180100756 to LBB), Flinders University and by the PhD scholarships received by AB from AJ and IM Naylon and the Playford Trust. We are sincerely grateful to those who assisted with the collection of samples, which included staff from the South Australian Research and Development Institute, Victorian Fisheries Authority and NSW Department of Primary Industries, as well as Tony Kemna (Far Out Charters), members of the Portland Sport Fishing Club and the organisers of the Kingston Offshore Fishing Competition 2019. We also sincerely thank Michelle Gardner for providing the temporal samples from Northern Gulf St Vincent and Port Phillip Bay. Lastly, we thank Andrea Barceló and Diana-Elena Vornicu at the Molecular Ecology Lab at Flinders University for their assistance in the laboratory.

## Supplementary data

Supplementary material is available at the ICESJMS online version of the manuscript.

## Author contributions

LBB, MW, AF, PH and JS conceptualized the study and acquired funding. LBB and AB conceived the idea for the manuscript. AB conducted the laboratory work, with contributions from JS and CB. AB analysed the data, with assistance from JS and CB. AB led the writing of the first version of the manuscript. All authors contributed to the revising of the final manuscript.

## Data availability statement

The SNP dataset is available on figshare: to be provided upon acceptance

