## Supplemental Files for "Bioregional boundaries and genomically-delineated stocks in snapper (*Chrysophrys auratus*) from southeastern Australia"

**Supplementary material**

**Table S1** Catch data for each snapper sample, including the two temporal ones from NGSV and PPB.

|  | Jurisdiction | Management area | Avg. lat | Avg. lon | Catch dates (mm/yy) | Sector |
| --- | --- | --- | --- | --- | --- | --- |
| Ceduna (CED) | South Australia | Spencer Gulf/  West Coast | -32.3 | 133.8 | 09/18, 08/19 | Commercial |
| Northern Spencer Gulf (NSG) | South Australia | Spencer Gulf/  West Coast | -33.2 | 137.7 | 09,10,12/18 | Commercial/  Research |
| Southern Spencer Gulf (SSG) | South Australia | Spencer Gulf/  West Coast | -34.3 | 136.6 | 08,10,12/18, 10/19 | Commercial/  Research |
| Northern Gulf St Vincent (NGSV) | South Australia | Gulf St Vincent | -34.5 | 138.1 | 08,10,11,12/18, 05/19 | Commercial/  Research |
| Northern Gulf St Vincent 2010 (NGSV10) | South Australia | Gulf St Vincent | -34.8 | 138.2 | 02/10 | Commercial |
| Southern Gulf St Vincent (SGSV) | South Australia | Gulf St Vincent | -35.3 | 138.2 | 07,08,10/18, 05,08/19 | Commercial |
| Kingston SE (KSE) | South Australia | Western Victoria | -36.5 | 139.4 | 04/19 | Recreational |
| Portland (PLD) | Victoria | Western Victoria | -38.4 | 142.0 | 01,02,05,06/19 | Recreational |
| Port Phillip Bay (PPB) | Victoria | Western Victoria | -38.0 | 144.9 | 11,12/18, 01,02/19 | Commercial |
| Port Phillip Bay 2011 (PPB11) | Victoria | Western Victoria | -38.2 | 144.8 | 2011 | Recreational |
| Western Port Bay (WPB) | Victoria | Western Victoria | -38.3 | 145.3 | 11/18 | Recreational |
| Lakes Entrance (LE) | Victoria | Eastern Victoria | -38.1 | 148.1 | 11/18 | Recreational |
| Eden (EDN) | New South Wales | East Coast | -37.1 | 150.1 | 04,05,10,11/18, 01/19 | Commercial |

**Table S2** Numbers of SNPs retained after each bioinformatics filtering step.

| Step | SNP count |
| --- | --- |
| Raw SNP catalogue | 7,342,804 |
| 80% of individuals, biallelic, >0.03 minor allele frequency | 50,142 |
| Remove indels | 45,940 |
| Read quality (ratio quality/coverage depth >0.2) | 45,362 |
| Mapping quality (>30) | 41,468 |
| High coverage loci (≤mean depth + (2*standard deviation)) | 40,419 |
| Hardy-Weinberg equilibrium in >67% of locations | 35,702 |
| Call error rate (0.95) | 32,221 |
| Linkage disequilibrium (500) | 11,266 |
| Putatively neutral loci | 10,916 |

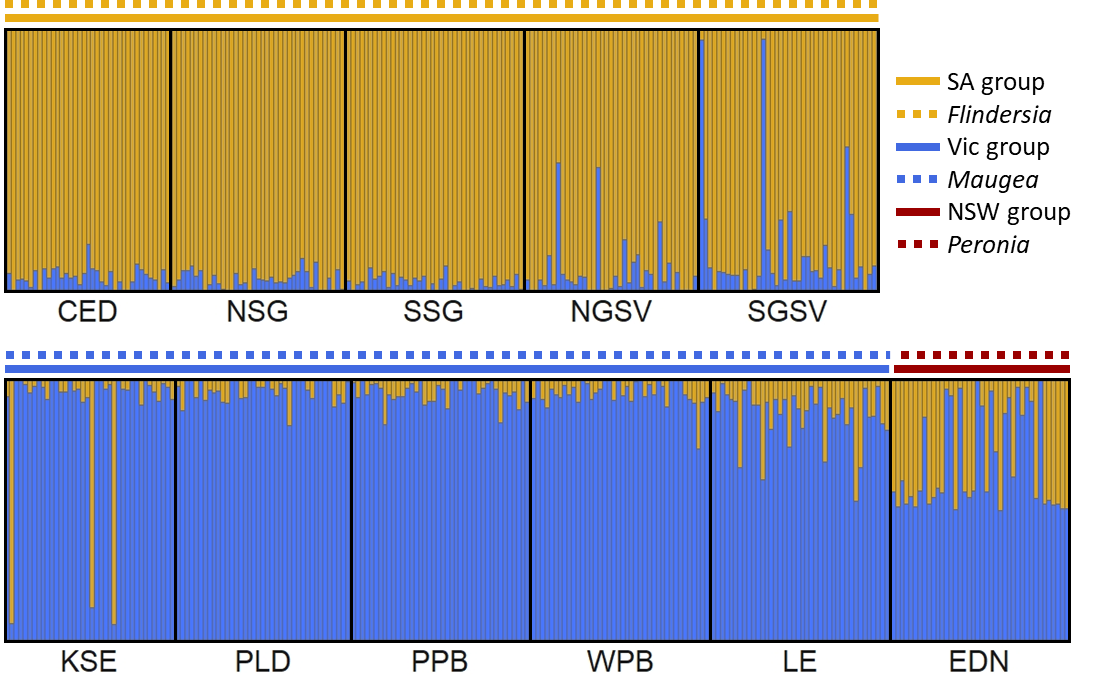

**Figure S1** ADMIXTURE results for *K* = 2 based on the 432 individuals (excluding the temporal samples from NGSV and PPB) and 10,916 neutral SNPs. Biogeographic provinces (Flindersia, Maugea, Peronia) and genetic groupings identified with ADMIXTURE (*K* = 3) and PCA (i.e., SA, Vic, NSW groups) are marked above the plots. Each vertical bar represents an individual and its probability of membership to each of the *K* groups is indicated by its colour makeup.

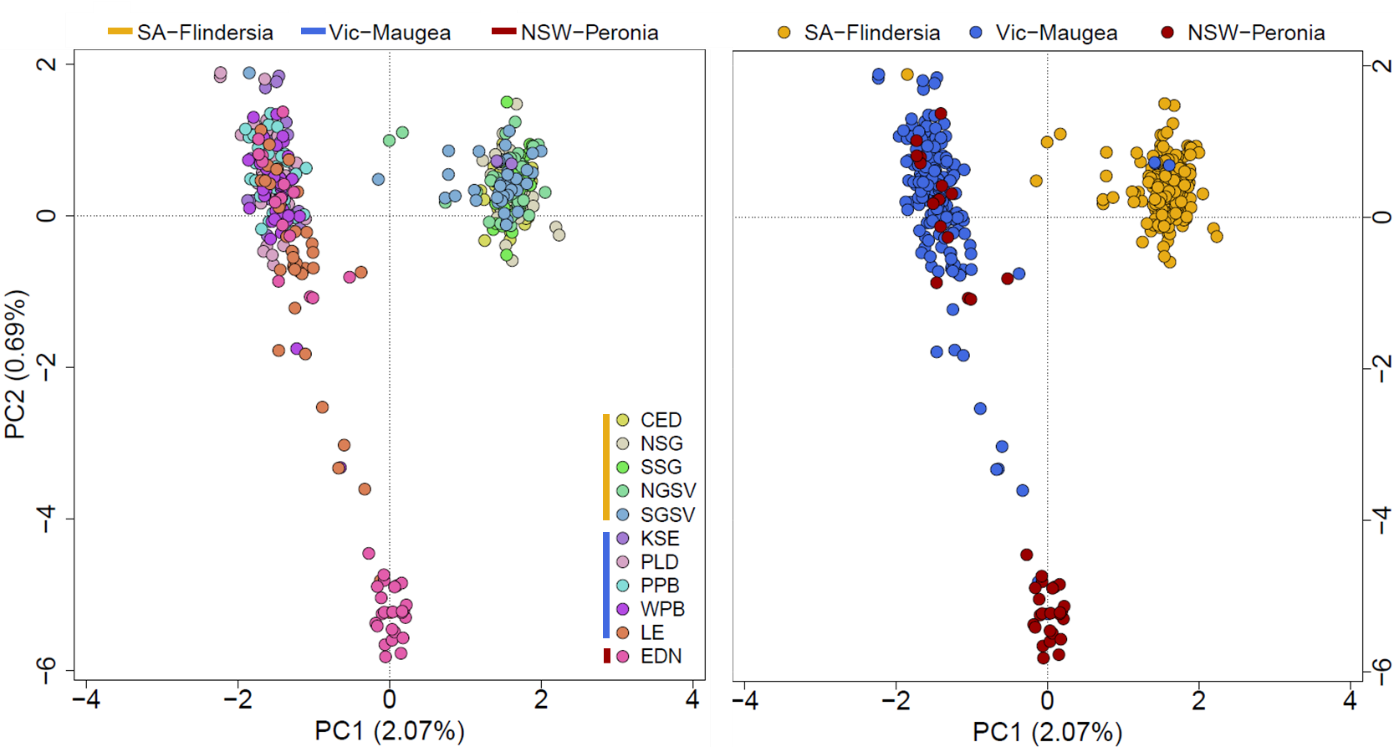

**Figure S2** Principal component analysis based on 10,916 neutral SNPs and the 432 individuals (excluding the temporal samples from NGSV and PPB), with individuals coloured by site in the left plot and by bioregion/genetic group in the right plot. The first two principal components are shown, and the proportion of genetic variance explained by each component is in parentheses.

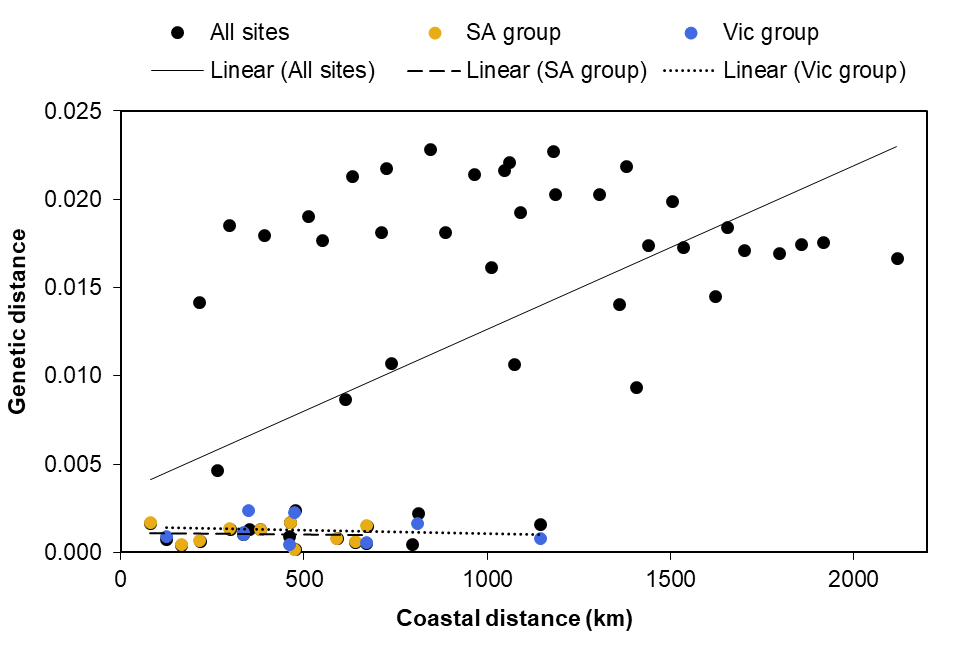

**Figure S3** Relationship between coastal distance and genetic distance (linearized *F*_ST_, *F*_ST_ /1- *F*_ST_) for all sites (black dots and solid trendline; r = 0.571), the SA group (gold dots and dashed trendline; r = 0.064) and the Vic group (blue dots and dotted trendline; r = 0.230). Only the Mantel test including all sites was significant (*p*-values = 0.003, 0.53 and 0.32 respectively).

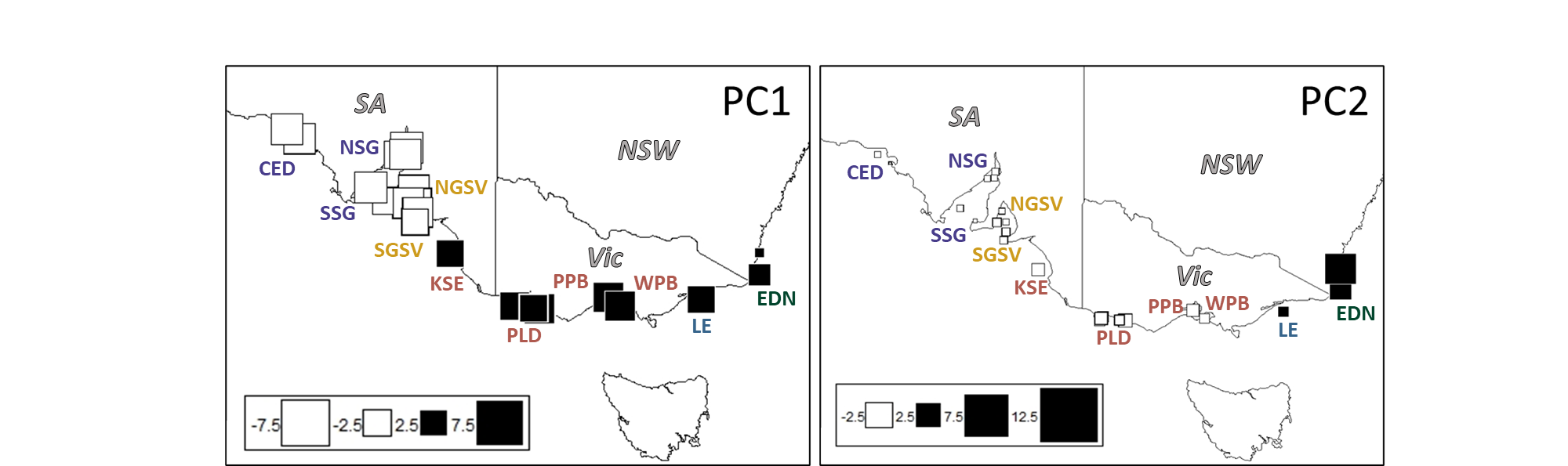

**Figure S4** The first and second global scores of spatial PCA based on all 11 snapper samples collected in 2018/9. Site names are coloured according to the management area in which they reside (as in Figure 1): ● = Spencer Gulf/West Coast, ● = Gulf St Vincent, ● = Western Victoria, ● = Eastern Victoria, ● = East Coast.
